# Semi-Supervised Learning of the Electronic Health Record for Phenotype Stratification

**DOI:** 10.1101/039800

**Authors:** Brett K. Beaulieu-Jones, Casey S. Greene, the Pooled Resource Open-Access ALS Clinical Trials Consortium

**Affiliations:** Graduate Group in Genomics and Computational Biology, Computational Genetics Lab, Institute for Biomedical Informatics, University of Pennsylvania; Department of Systems Pharmacology and Translational Therapeutics, Institute for Biomedical Informatics, Institute for Translational Medicine and Therapeutics, University of Pennsylvania

**Keywords:** Electronic Health Record, Denoising Autoencoder, Unsupervised, Electronic Phenotyping, Patient Stratification, Disease Subtyping

## Abstract

Patient interactions with health care providers result in entries to electronic health records (EHRs). EHRs were built for clinical and billing purposes but contain many data points about an individual. Mining these records provides opportunities to extract electronic phenotypes, which can be paired with genetic data to identify genes underlying common human diseases. This task remains challenging: high quality phenotyping is costly and requires physician review; many fields in the records are sparsely filled; and our definitions of diseases are continuing to improve over time. Here we develop and evaluate a semi-supervised learning method for EHR phenotype extraction using denoising autoencoders for phenotype stratification. By combining denoising autoencoders with random forests we find classification improvements across multiple simulation models and improved survival prediction in ALS clinical trial data. This is particularly evident in cases where only a small number of patients have high quality phenotypes, a common scenario in EHR-based research. Denoising autoencoders perform dimensionality reduction enabling visualization and clustering for the discovery of new subtypes of disease. This method represents a promising approach to clarify disease subtypes and improve genotype-phenotype association studies that leverage EHRs.

**GRAPHICAL ABSTRACT:** 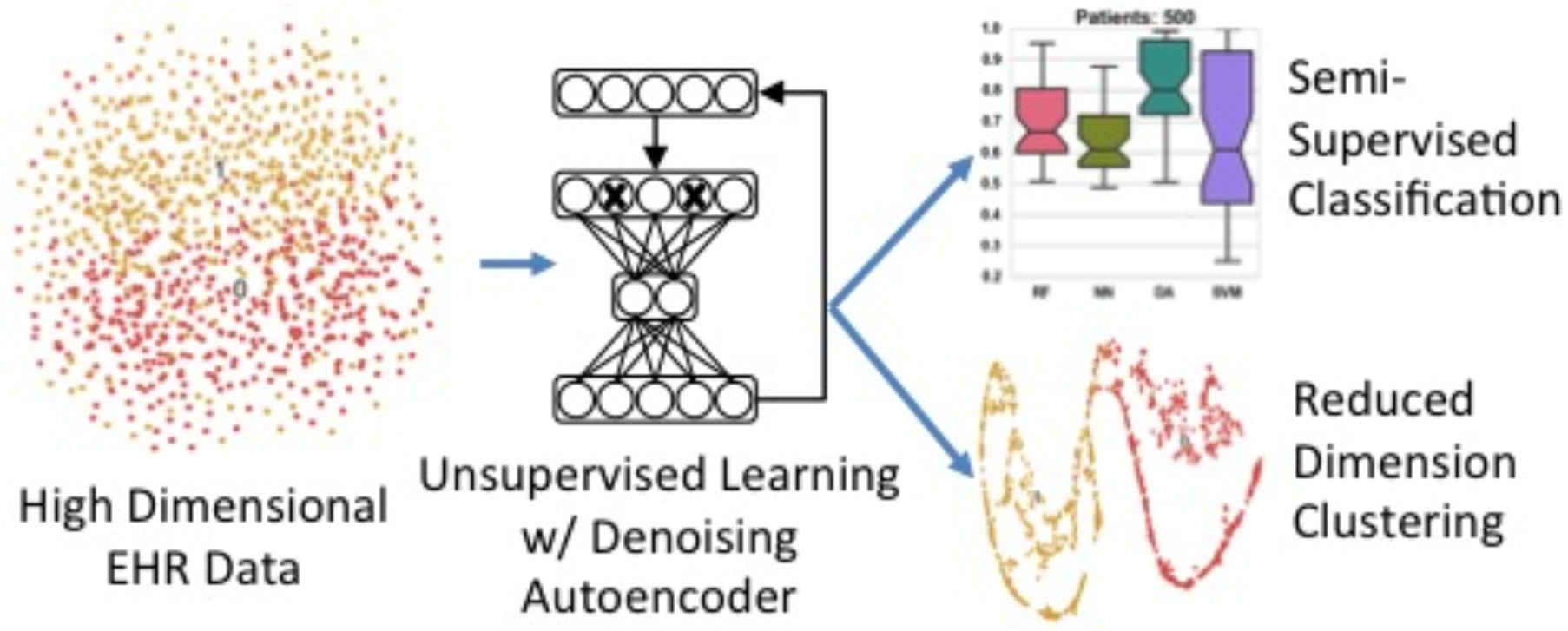

**HIGHLIGHTS:**

- Denoising autoencoders (DAs) can model electronic health records.
- Semi-supervised learning with DAs improves ALS patient survival predictions.
- DAs improve patient cluster visualization through dimensionality reduction.

## INTRODUCTION

Biomedical research often considers diseases as fixed phenotypes, but many have evolving definitions and are difficult to classify. The electronic health record (EHR) is a popular source for electronic phenotyping to augment traditional genetic association studies, but there is a relative scarcity of research quality annotated patients [1]. Electronic phenotyping relies on either codes designed for billing or time intensive manual clinician review. This is an ideal environment for semi-supervised algorithms, performing unsupervised learning on many patients followed by supervised learning on a smaller, annotated, subset. Denoising autoencoders (DAs) are a powerful tool to perform unsupervised learning [2]. DAs are a type of artificial neural network trained to reconstruct an original input from an intentionally corrupted input. Through this training they learn higher-level representations modeling the structure of the underlying data. We sought to determine whether applying DAs to the EHR could reduce the number of annotated patients required, construct non-billing code based phenotypes and elucidate disease subtypes for fine-tuned genetic association.

The United States federal government mandated meaningful use of EHRs by 2014 to improve patient care quality, secure and communicate patient information, and clarify patient billing [3,4]. Despite not being designed specifically for research, EHRs have already proven an effective source of phenotypes in genetic association studies [5,6]. Initially, phenotypes were hand designed based on manual clinician review of patient records. These studies were limited by the time and cost inherent in manual review, but DAs can make use of unlabeled data [7]. After unsupervised pre-training, the trained DA’s hidden layer can be used as input to a traditional classifier to create a semi-supervised learner. This allows the DA to learn from all samples, even those without labels, and requires only a small subset to be annotated. Today, phenome-wide association studies (PheWAS) are the most prevalent example of EHR phenotyping, proving particularly effective at identifying pleiotropic genetic variants [8]. PheWASs often use algorithms based on the International Classification of Disease (ICD) codes to construct a phenotype. This coding system was designed for billing, not to capture research phenotypes. DA constructed features are combinations of many components of clinical data and may provide a more holistic view of a patient than billing codes alone.

Through extensive study, disease diagnoses can become more precise over time [9–13]. Cancers, for example, were historically typed by occurrence location and the efficacy of different treatments. As the mechanisms of cancer are better understood, they are further categorized by their physiological nature. The progression of subtypes in lung cancer illustrates this increased understanding over time [9]. Beginning with a single diagnosis based on occurrence in the lung, lung cancer has been divided into dozens of subtypes over several decades based on histological analysis, and genetic markers [10–13]. The unsupervised nature of DAs means that even if the definitions of a disease change, they would not need to be retrained. The ability to identify more homogenous phenotypes showed increased genotype to phenotype linkage in schizophrenia, bipolar disease [14], and Rett Syndrome [15–18]. Furthermore, type 2 diabetes subtypes have been discovered using topological analysis of EHR patient similarity [19]. The dimensionality reduction possible with a DA makes clustering and visualization more feasible. Subtyping exposes disease heterogeneity and may contribute to additional physiological understanding.

Previous work in semi-supervised learning of the EHR relies on closed source commercial software [19], and natural language processing of free text fields to match clinical diagnosis [20,21]. We are not aware of any previous work performing semi-supervised classification and clustering from quantitative structured patient data.

We evaluate DAs for phenotype construction using four simulation models of EHR data for complex phenotypes, modify DAs to effectively handle missingness in data and use the DA to create cluster visualizations that can aid in the discovery of subtypes of complex diseases. We apply these methods to predict ALS patient survival and to visualize ALS patient clusters. ALS is a progressive neurodegenerative disorder, which attacks the neurons responsible for controlling muscle function [22]. ALS patients typically die within 3 to 5 years, but some patients can survive more than 10 years, the disease is considered clinically heterogeneous and predicting the rate of progression can be challenging [23].

## METHODS

We developed an approach, entitled “Denoising Autoencoders for Phenotype Stratification (DAPS),” that constructs phenotypes through unsupervised learning. This generalized phenotype construction can be used to classify whether patients have a particular disease or to search for disease subtypes in patient populations. To evaluate DAPS, we created a simulation framework with multiple hidden factors influencing potentially overlapping observed variables. We evaluated the reduced DA models against feature-complete representations with popular supervised learning algorithms. These evaluations covered both complete datasets, as well as the more realistic cases of incompletely labeled and missing data. We developed a technique that uses the reduced feature-space of the DA to visualize potential subtypes. Finally, we evaluate DAs ability to predict ALS patient survival in both classification and clustering tasks. Each of these is fully described below and full parameters included in sweeps are available in the supplementary materials.

Source code to reproduce each analysis is included in our repository (https://github.com/greenelab/DAPS) [24] and is provided under a permissive open source license (3-clause BSD). A docker build is included with the repository to provide a common environment to easily reproduce results without installing dependencies [25]. In addition, Shippable, a continuous integration platform, is used to reanalyze results in a clean environment and generate figures after each commit [26].

### Unsupervised Training with Denoising Autoencoders

DAs were initially introduced as a component in constructing the deep networks used in deep learning [27]. Deep learning algorithms have become the dominant performers in many domains including image recognition, speech recognition and natural language processing [28–33]. Recently they have also been used to solve biological problems including tumor classification, predicting chromatin structure and protein binding [2,34,35]. DAs showed strong performance early in the deep learning revolution but have been surpassed in most domains by convolutional neural networks or recurrent neural networks [27]. While these complex deep networks have surpassed the performance of DAs in these areas, they rely on strictly structured relationships such as the relative positions of pixels within an image [30,36]. This structure is unlikely to exist in the EHR. In addition, complex deep networks are notoriously hard to interpret. DAs are easily generalizable, benefit from both linear and nonlinear correlation structure in the data, and contain accessible, interpretable, internal nodes [2]. Oftentimes the hidden layer is a “bottle-neck”, a much smaller size than the input layer, in order to force the autoencoder to learn the most important patterns in the data [36].

We used the Theano library [37,38] to construct a DA consisting of three layers, an input layer *x*, a single hidden layer *y*, and a reconstructed layer *z* [27] (Figure 1A). Noise was added to the input layer through a stochastic corruption process, which masks 20% of the input values, selected at random, to zero.

**Fig. 1.**
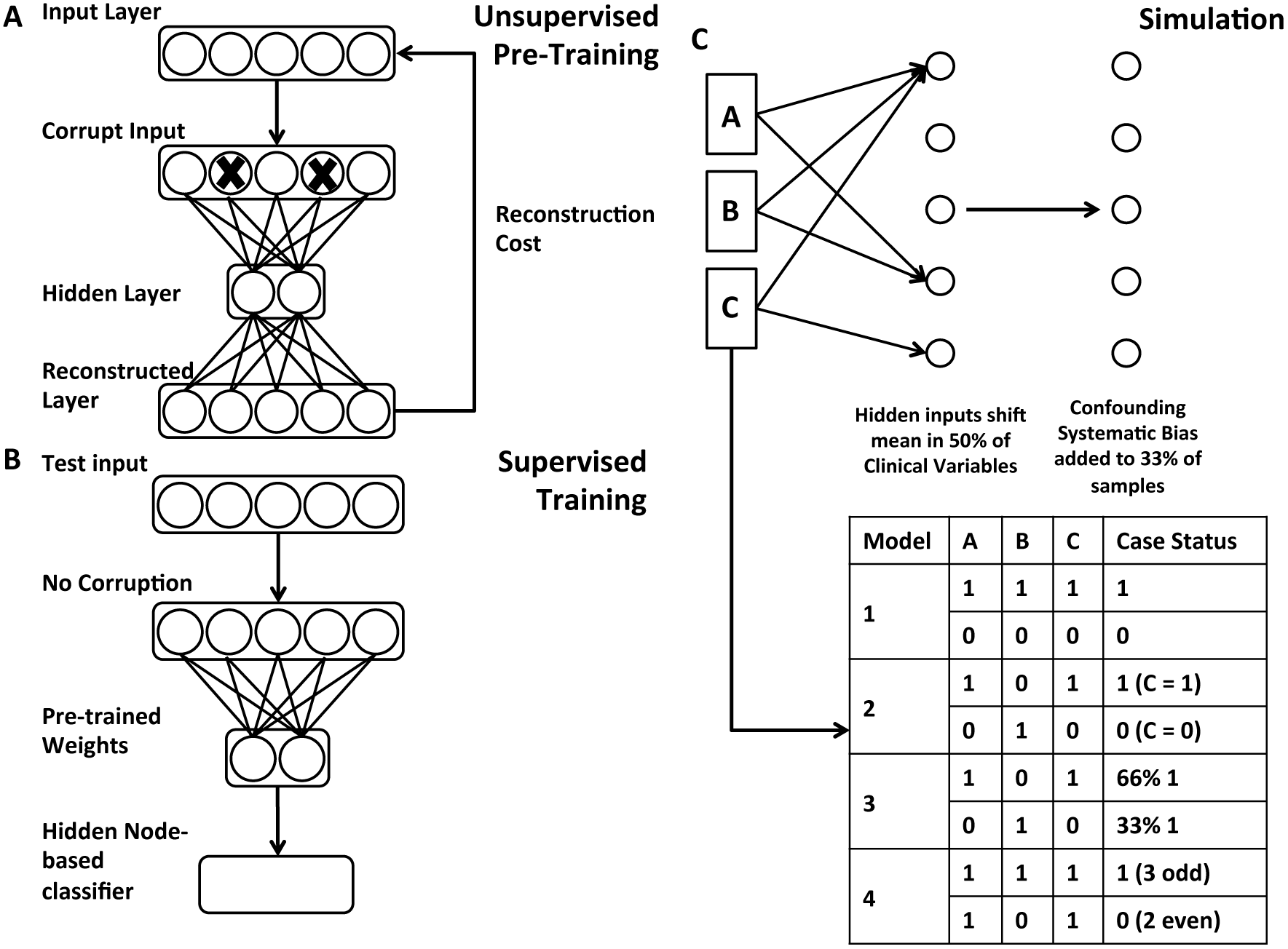
**A)** Network diagram of DAs used for unsupervised pre-training. Input data is intentionally corrupted and then weights and biases are learned to minimize reconstruction cost when mapping the data to a hidden layer and back to a reconstructed layer. **B)** Supervised classification occurs using the pre-trained DA hidden nodes as input to a traditional classifier. **C)** Simulation model with example cases and controls under each rule set.

The hidden layer y was calculated by multiplying the input layer by a weight vector *W*, adding a bias vector *b* and computing the sigmoid (Formula 1). The reconstructed layer *z* was similarly computed using tied weights, the transpose of *W*and *b* (Formula 2). The cost function is the cross-entropy of the reconstruction, a measure of distance between the reconstructed layer and the input layer (Formula 3).

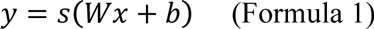

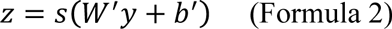

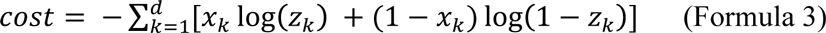

Stochastic gradient descent was performed for 1000 training epochs, at a learning rate of 0.1. Hidden layers of two, four, eight and sixteen hidden nodes were included in the parameter sweep with a 20% input corruption level. Vincent et al. [27] provide a through explanation of training for DAs without missing data.

In the event of missing data, the cost calculation was modified to exclude missing data from contributing to the reconstruction cost. A missingness vector *m* was created for each input vector, with a value of 1 where the data is present and 0 when the data is missing. Both the input sample *x* and reconstruction *z* were multiplied by *m* and the cross entropy error was divided by the sum of the *m*, the number of non-missing features to get the average cost per feature present (Formula 4). This allowed the DA to learn the structure of the data from present features rather than imputation.

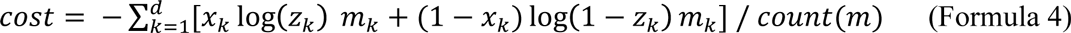

### Supervised Denoising Autoencoder Classifier

To convert the DA to a supervised classifier, we first trained the DA in an unsupervised fashion (pre-training) (Fig 1A). We then applied a variety of traditional machine learning classifiers including, decision trees, random forests, logistic regression, nearest neighbors and support vector machines to the pre-trained unsupervised hidden layer values, *y*, of the DA (Figure 1B). Random forests applied to DA hidden nodes (DA+RF) were shown for all comparisons. Predictive performance was measured by comparing the AUROC using stratified 10-fold cross validation. The Scikit-learn library was used for the traditional classifiers [39]. The Support Vector Machine uses a radial basis function kernel, with a penalty parameter of 1. The nearest neighbors classifier uses a k-value of 5 and the random forest uses 10 estimators. These parameters achieved optimal performance in a preliminary parameter sweep.

### Simulation Framework

We designed four simulation models to evaluate algorithmic performance. These simulations were not designed to perfectly recapitulate EHR data. Instead they are designed to capture a variety of complexity in order to identify algorithmic strengths and weaknesses with known underlying models.

To simulate patients, first clinical observations were generated by first drawing random samples from a normal distribution. Next hidden input effects were generated in accordance with one of four simulation models. When turned on these hidden input effects shift 1 to N observed clinical variables with replacement (Figure 1C). Shifted clinical features were chosen at random, but consistent for all patients. Case-control status was determined by rules applied to the hidden input effects, where 1 represents the effect being on and 0 represents the effect being off. Next, a confounding systematic bias was added to a random subset (33%) of the patients as a source of additional noise to simulate the variance accompanying data created by physicians, labs, hospitals or other spurious effects.

There are four models defining hidden input effect rules to determine case-control status:

1. **All together/all relevant.** Individuals have the same value (0 or 1) for all hidden input effects. Controls have all hidden effects set to 0. Cases have all hidden effects set to 1. A model capturing any hidden input will be able to predict case/control status in this scenario. This is a test of whether each method can recognize any of the hidden effects.
2. **All independent /single effect relevant.** Individuals have 0 to N (specified per simulation) hidden input effects chosen at random. One arbitrary effect (the last one) is used to determine case-control status. In controls, this is 0. In cases, this is 1. A model capturing the relevant hidden input will be able to predict case/control status in this scenario. This is a test of whether each method can recognize the important hidden effect when there are multiple shifted distributions.
3. **All independent/percentage based.** Individuals have 0 to N (specified per simulation) of hidden input effects chosen at random set to 1. The percentage of hidden input effects on represents the probability of the patient being a case. A model capturing more hidden effects will be able to more accurately predict case/control in this scenario. This is a test of whether each method can perform effectively without a hard-rule based model, and could represent a disease with incomplete penetrance.
4. **All independent/complex rule based.** Individuals have 0 to N (specified per simulation) of hidden inputs chosen at random set to 1. The sum of hidden effects determines case-control status (cases are even, controls are odd). A model must capture all hidden effects to successfully predict case/control in this scenario. This is a test to identify the complexity limitations of each of the methods.

Simulation model 3 could reflect a disease with incomplete penetrance. The probabilistic manner of this simulation means that the optimum binary classifier will have an expected accuracy limited by the role of stochasticity in the model. The amount of stochasticity is a function of the number of hidden effects. In this model, case control odds were equal to the percentage of hidden input effects on. If there are 4 hidden input effects and 2 are on, the patient has a 50% chance of being a case and a 50% chance of being a control. A binary classifier can perfectly model this simulation and still have error due to the probabilistic nature. The maximum expected accuracy was calculated from a binomial distribution multiplied by the minority percentage as the best a binary classifier could do is choose the more likely class. For example, in the case of 4 hidden effects the maximum expected accuracy is 68.75%.

An example of a hypothetical condition a hidden input effect could represent is the familial hypercholesterolemia genotype. For a patient with the familial hypercholesterolemia genotype, the simulated clinical observations could represent increases in levels of total and low-density lipoprotein cholesterol, the deposition of cholesterol in extravascular tissues, corneal arcus and elevated triglyceride levels [40]. Some factors such as elevated triglyceride levels are not solely the result of the genetic predisposition and are related to environmental factors. Hypothetically additional hidden input effects on the same observed variable would represent these other factors. Because our goal is to evaluate methods for their ability to broadly capture these types of patterns, we generate randomized relationships between hidden and observed variables. This avoids overfitting our evaluation to specific phenotypes.

#### Supervised Classification Comparison

If successfully trained, the hidden layer of a DA, *y*, captures the first *n* factors of variation in the data, where *n* is the number of nodes in the hidden layer. To test whether the DA constructed useful features by learning the main factors of variation in the data we used the trained hidden layer as an input to a shallow classifier.

To do this, we first completed unsupervised pre-training of the DA with all of the simulated samples. The hidden layer values, *y*, were calculated for all samples using the trained DA without any corruption and fed in as the features to a random forest to form a supervised classifier.

Classification performance between DAs plus random forests (DA+RF) were compared against decision trees, random forests, nearest neighbors and support vector machines in a parameter sweep under each model (Table 1). Additional model parameters included in sweeps are included in the supplementary materials. All traditional classifiers were implemented with Scikit-learn [39]. Classification performance was compared using the AUROC with 10-fold cross validation across 10 independent replicates for each set of parameters.

**Table 1.**
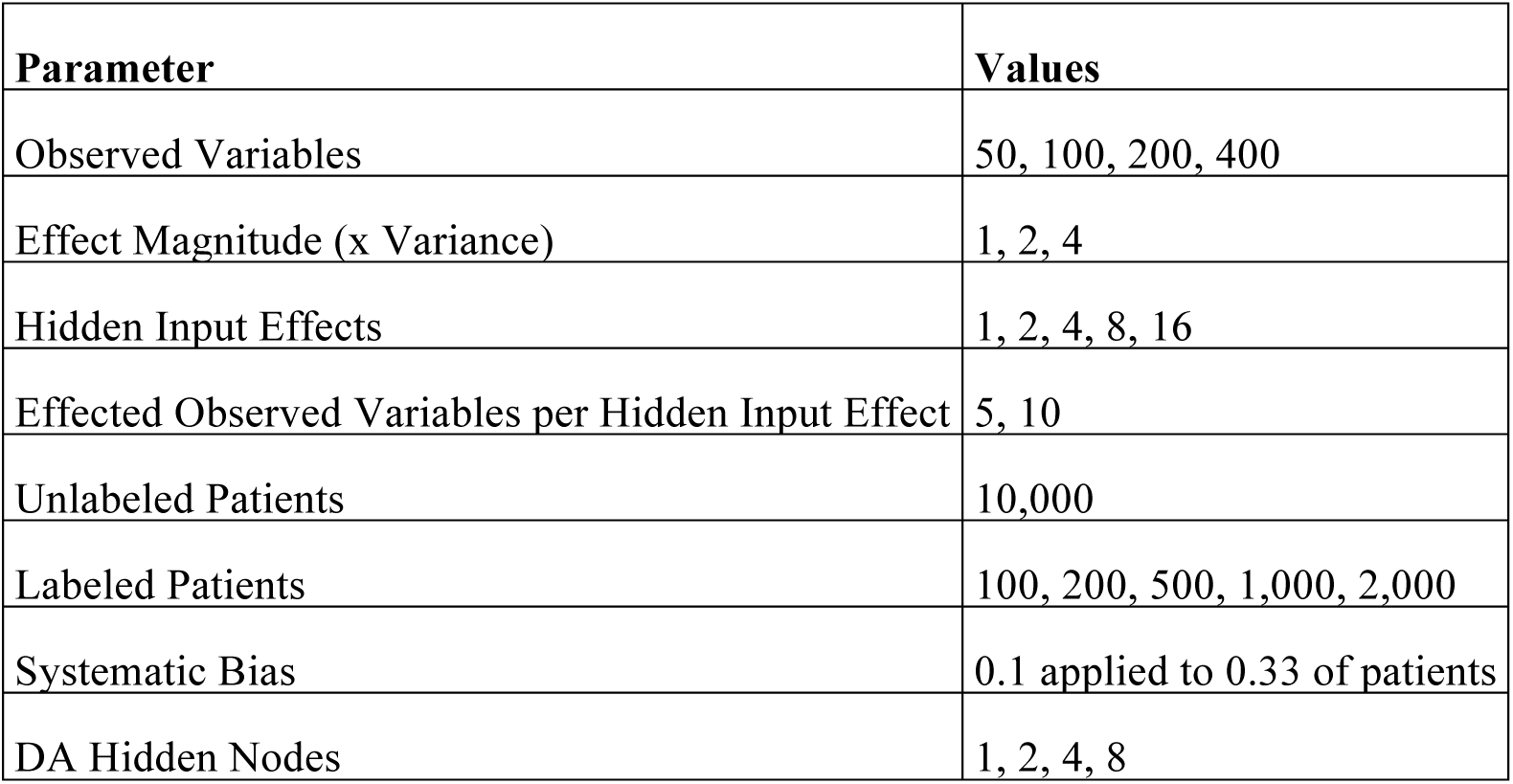
Simulation Model 1 Parameter Sweep Specifications.

#### Semi-Supervised Classification Comparison

The supervised classification comparison was repeated but with additional patients simulated and utilized during the unsupervised pre-training of the DA. The additional patients were simulated at the same 50% case, 50% control ratio but their labels were discarded after simulation. These additional patients were mixed with the original labeled patients and included in the unsupervised pre-training of the DA. The unlabeled samples were then discarded and the DA+RF was then provided the same, labeled, patient groups as the traditional classifiers. The labeled patient samples were run through the trained DA in the same manner as the unsupervised pre-training but without any corruption added to the data. The DA+RF and traditional classifiers were evaluated in a parameter sweep under each model using 10-fold cross validation.

#### Missing Data Comparison

The semi-supervised classification comparison was repeated five times with, 0%, 10%, 20%, 30% and 40% of the data missing. Missing data was added at random per sample, depending on the specified percentage missing.

Throughout these trials, the cost calculation was modified to exclude missing data from the cost and allow the DA to learn without imputing values (Formula 4). The traditional classifiers were trained using mean imputation for missing data. Mean imputation is particularly well suited for the simulation models because the observations were drawn from normal distributions, potentially giving an advantage to the non-DA algorithms that would not be available in many real datasets.

As in the semi-supervised classification comparison trial, the DA+RF and traditional classifiers were evaluated under each model using 10-fold cross validation.

#### Clustering and Visualization

To interpret and visualize results, patient populations were clustered using principal components analysis (PCA) and t-Stochastic Neighbor Embedding (t-SNE) of the trained DA’s hidden nodes [41,42]. PCA and t-SNE were implemented with the Sci-kit learn library [39].

Ten thousand patients (5,000 cases, 5,000 controls) with four hidden effects were simulated under model 1. PCA followed by t-SNE was performed initially on the raw input for comparison and then on the hidden nodes of the DA after every 10 training epochs.

To test the ability to identify subtypes, we simulated 15,000 patients, 5,000 cases under model 1, 5,000 cases under model 2, and 5,000 controls. Input observations were compared to two, three and four-node DAs using PCA followed by t-SNE.

### ALS Survival Analysis

Data used in the ALS Survival portion of this article were obtained from the Pooled Resource Open-Access ALS Clinical Trials (PRO-ACT) Database. In 2011, Prize4Life, in collaboration with the Northeast ALS Consortium, and with funding from the ALS Therapy Alliance, formed the Pooled Resource Open-Access ALS Clinical Trials (PROACT) Consortium. The data available in the PRO-ACT Database has been volunteered by PRO-ACT Consortium members. The PRO-ACT dataset includes 23 clinical trials covering 10,723 patients. We limit our survival analysis to the 3,398 patients with known death information, but perform unsupervised pre-training of the DA with all 10,723 patients.

Patient data includes quantitative features consisting of demographic information, diagnosis history, family history, treatment history, vital sign readings, concomitant medications and laboratory tests. Categorical variables were converted to one-hot encoding. Repeated or temporal measurements were encoded as the mean, minimum, maximum, count, standard deviation and slope across each repeat. Measurement scales were standardized and input features were normalized to be between 0 and 1. No imputation was performed on the input to the DA; K-nearest neighbors imputation (K=15) was performed for the raw comparison. This preprocessing resulted in an input layer of 6,812 numerical features per patient.

Patient survival was predicted as the number of days from disease onset. Random Forest Regression using Scikit-learn with 1,000 estimators was performed on the raw data and the hidden layer of a 250 node DA trained for 1,000 epochs. Performance was evaluated using 10-fold cross validation. Cluster analysis using t-SNE was compared between PCA (2, 4, 8, and 16 components) on the raw input with the hidden layer of the DA.

## RESULTS

### Case-Control DA Training Visualization

We trained a DA and visualized the training process using PCA and t-SNE. These visualization techniques offer intuition and the ability to examine the sub-clusters. Given 5,000 cases and 5,000 controls under simulation model 1, PCA and t-SNE alone did not yield defined clusters (Fig 2A). Figures 2B-F show the separation of cases from controls as the DA is trained. One thousand epochs of training via stochastic gradient descent were found to be sufficient for the convergence of reconstruction cost and stabilization of visualizations within simulated data (Figure 2 E and F, Supplemental Figure 1).

**Fig. 2.**
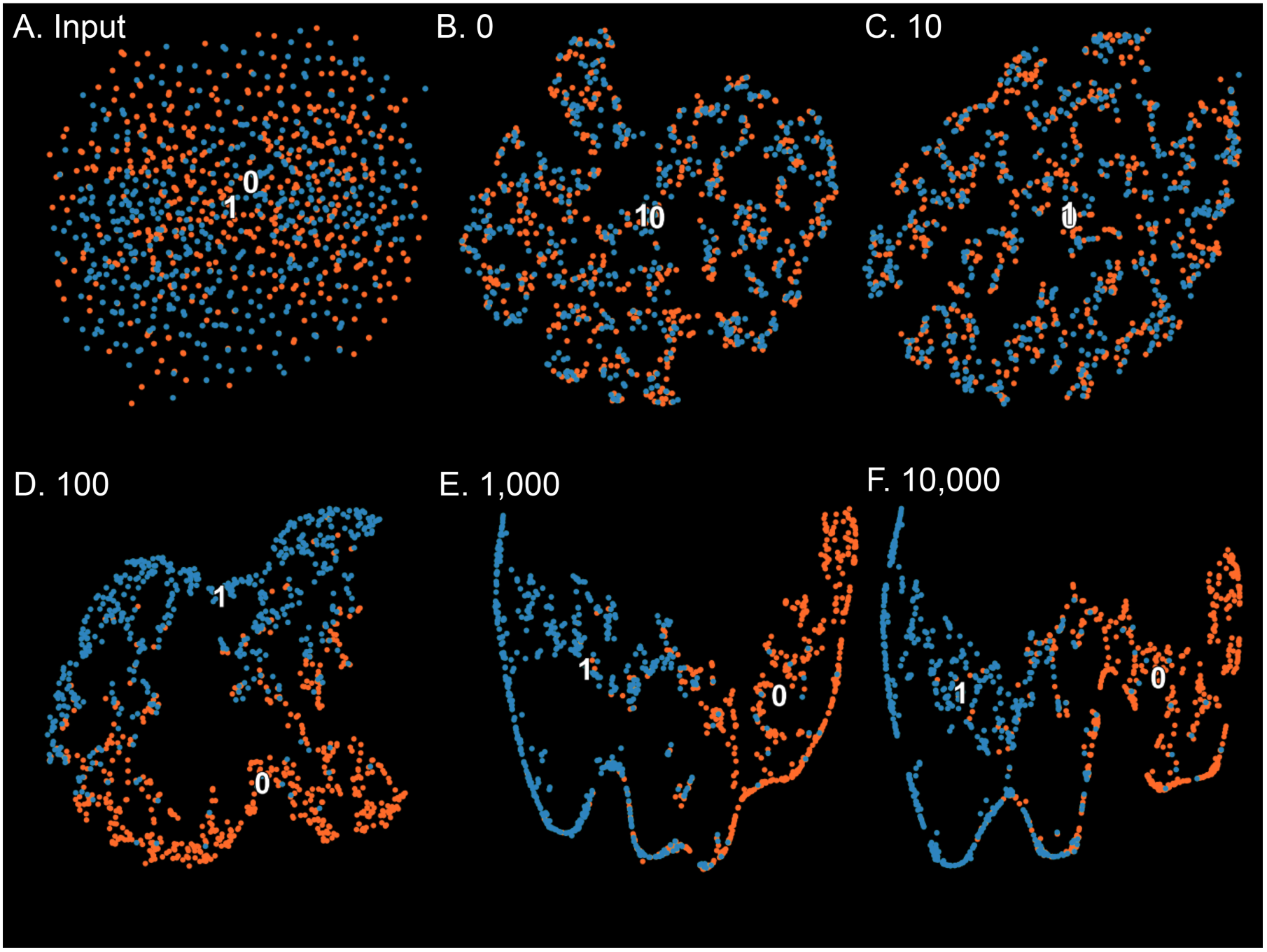
Case vs. Control clustering via principal components analysis and t-distributed stochastic neighbor embedding throughout the training of the DA. Controls are shown in yellow, cases are shown in red. **A.)** Raw input **B.)** 0 training epochs **C.)** 10 training epochs **D.)** 100 training epochs **E.)** 1,000 training epochs **F.)** 10,000 training epochs.

### Fully Supervised Comparison

To examine the ability of DAs to learn the structure of the data we compare the predictive ability of classification algorithms applied to the DA constructed through unsupervised training. Random forests demonstrated a strong balance of performance and stability, and were used for all comparisons (Supplemental Figure 2). We then compare the DA plus a random forest classifier (DA+RF) to the top performing classifiers on raw input data (Table 2).

**Table 2.** Mean Receiver Operating Curve Area Under Curve by method under simulation model 1. (10 Replicate, 10-fold cross validation)

| **Patients** | **DA+RF** | **Random Forest** | **Support Vector Machine** | **Decision Tree** | **Nearest Neighbors** |
| --- | --- | --- | --- | --- | --- |
| **100** | 0.618 | 0.653 | 0.504 | 0.599 | 0.635 |
| **200** | 0.637 | 0.610 | 0.449 | 0.589 | 0.608 |
| **500** | 0.677 | 0.690 | 0.663 | 0.617 | 0.642 |
| **1000** | 0.774 | 0.717 | 0.776 | 0.634 | 0.651 |
| **2000** | 0.755 | 0.736 | 0.862 | 0.643 | 0.658 |
| **Mean** | 0.692 | 0.681 | 0.651 | 0.616 | 0.639 |

Key trends emerged under each model; with few patients SVMs had AUCs indistinguishable from those expected from a random classifier. As one would expect, SVMs were top performers at when the number of patients was high. Random forest classification performance scaled steadily with patient count. The DA+RF performed similarly to the random forest, showing that a 2-node DA is able to capture at least one of the input hidden effects. Capturing any signal is sufficient to accurately classify simulation model 1.

### Semi-Supervised Comparison

The full potential of the DA+RF is reflected in semi-supervised parameter sweep comparison for simulation model 1 (Fig 3A). Each set of parameters was evaluated with 10 replicates and 10-fold cross validation for each replicate. With sufficient unlabeled examples, the DA method’s performance is high, even with very few labeled examples. Because of the extreme feature reduction, the traditional classifier on top of the DA is able to reach its learning capacity with very few labeled patients (Fig 3A). Efficient learning from labeled examples is critical in practical use cases because there are often few well-annotated cases due to the expense of clinician manual review. The 2-node DA plus random forest also showed strong performance in relation to an SVM when there were many observed clinical variables (Supp. Fig 3) and when there were many hidden effects. The SVM again showed the highest performance at very high numbers (1000 or more) of labeled patients. The advantages of semi-supervised learning diminish as the number of labeled patients gets closer to the number of total patients. In addition, at high patient counts, a DA with more than 2 hidden nodes is required to capture the structure of the data with higher resolution.

**Fig. 3.**
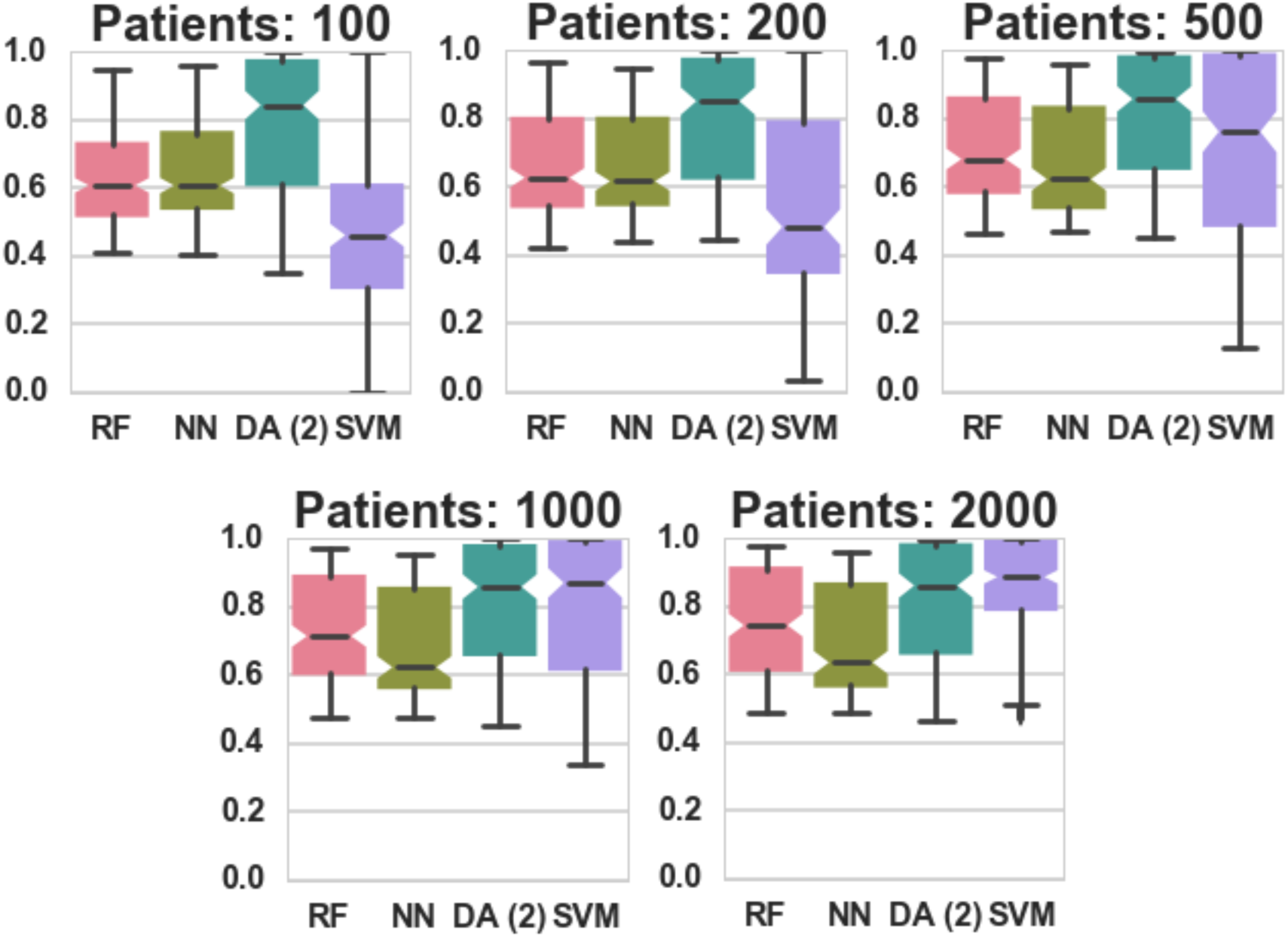
Classification AUC in relation to the number of labeled patients under simulation model 1 (RF – Random Forest, NN – Nearest Neighbors, DA – 2-node DA + Random Forest, SVM – Support Vector Machine). Unsupervised pre-training of the 2-node DA was performed with 10,000 patients. Notch indicates 95% confidence interval for the median. Whiskers extend 1.5 times past the low and high quartiles. Points outside this range are denoted as dots and represent outliers.

These patterns repeat across the other simulation models, with more complex models requiring more hidden nodes to adequately model the structure of the data. In simulation model 2 (Fig 4A), both 4 and 8 node DAs outperform the 2-node DA. Under Model 3, the 4-node DA is the strongest performing, with median performance 5% better than the next best traditional classifier. The 4-node DA’s median 95% confidence interval was above any of the compared methods. Model 4 (Fig 4C, 4D) was the most difficult to classify as the classifier had to capture all of the hidden effects to be accurate. In several cases, no classifier did better than the expected performance of a random classifier. In fact, the SVM’s average AUC over the entire sweep was statistically indistinguishable from random performance. As expected, the 2-node performs worse than the 4 and 8-node DAs on model 4. The 2-node DA lacks sufficient dimensionality to capture more than 4 hidden input effects.

**Fig. 4.**
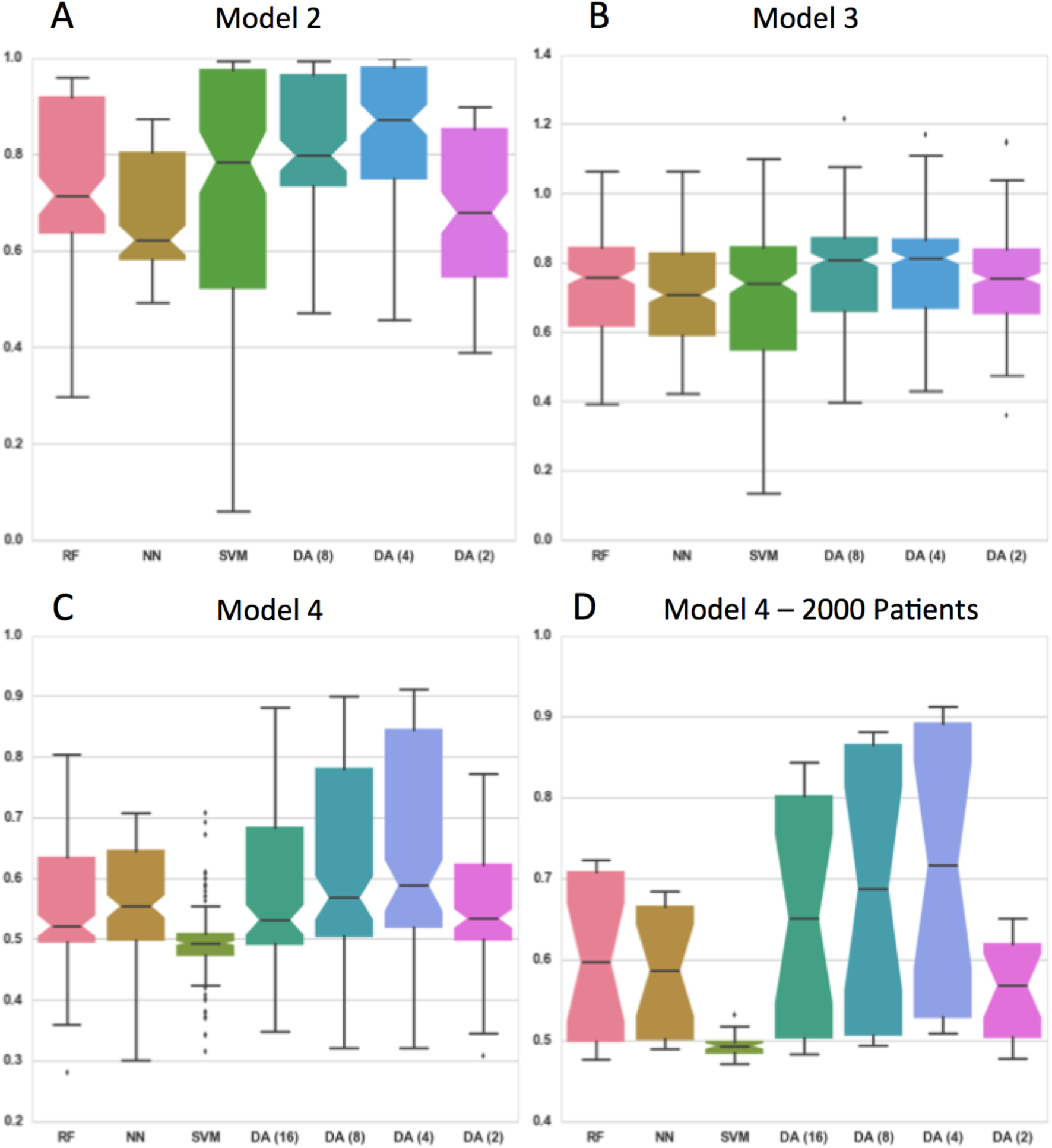
Classification accuracy comparisons for models 2-4. Notch indicates 95% confidence interval for the median. Whiskers extend 1.5 times past the low and high quartiles. **A)** Classification Accuracy of model 2 (1, 2, 4 and 8 effects). **B)** Classification AUC normalized to simulation model 2 expected max predictive accuracy (1, 2, 4 and 8 effects). **C)** Classification AUC of model 4 (1, 2, 4 and 8 effects). **D.)** Classification AUC of model 4 (parameter sweep results for 1, 2, 4 and 8 effects using only the parameter sets with 2,000 labeled patients)

Clinical records often have empty fields, so algorithms must be robust to missing data. We evaluated the DA’s robustness in this situation. The DA is robust to missing data maintaining near-max classification performance across the missingness proportions tested (Supp. Fig 4). For these simulation models, the mean imputation used for non-DA approaches is an ideal strategy. DAs and SVMs show consistent performance even as the percent of data missing increases, suggesting that the DA is at least as robust as the ideal imputation method.

### Simulated Subtype Clustering Visualization

We evaluated the DAPS’ ability to cluster patients for subtype identification. To perform this analysis, we simulated 5,000 cases from each of two different models (1 and 2) to represent a disease with two subtypes. An additional 5,000 controls were simulated. We then visualized the DA constructed from this set of patients using PCA followed by t-SNE. In the input data, the subtypes are relatively overlapping (Fig 5A). A DA with two nodes was also unable to separate this number of subtypes (Figure 5B). Visualizations constructed from DAs with three (Figure 5C) or four (Figure 5D) nodes were able to effectively separate both subtypes of cases from each other and from controls.

**Fig. 5.**
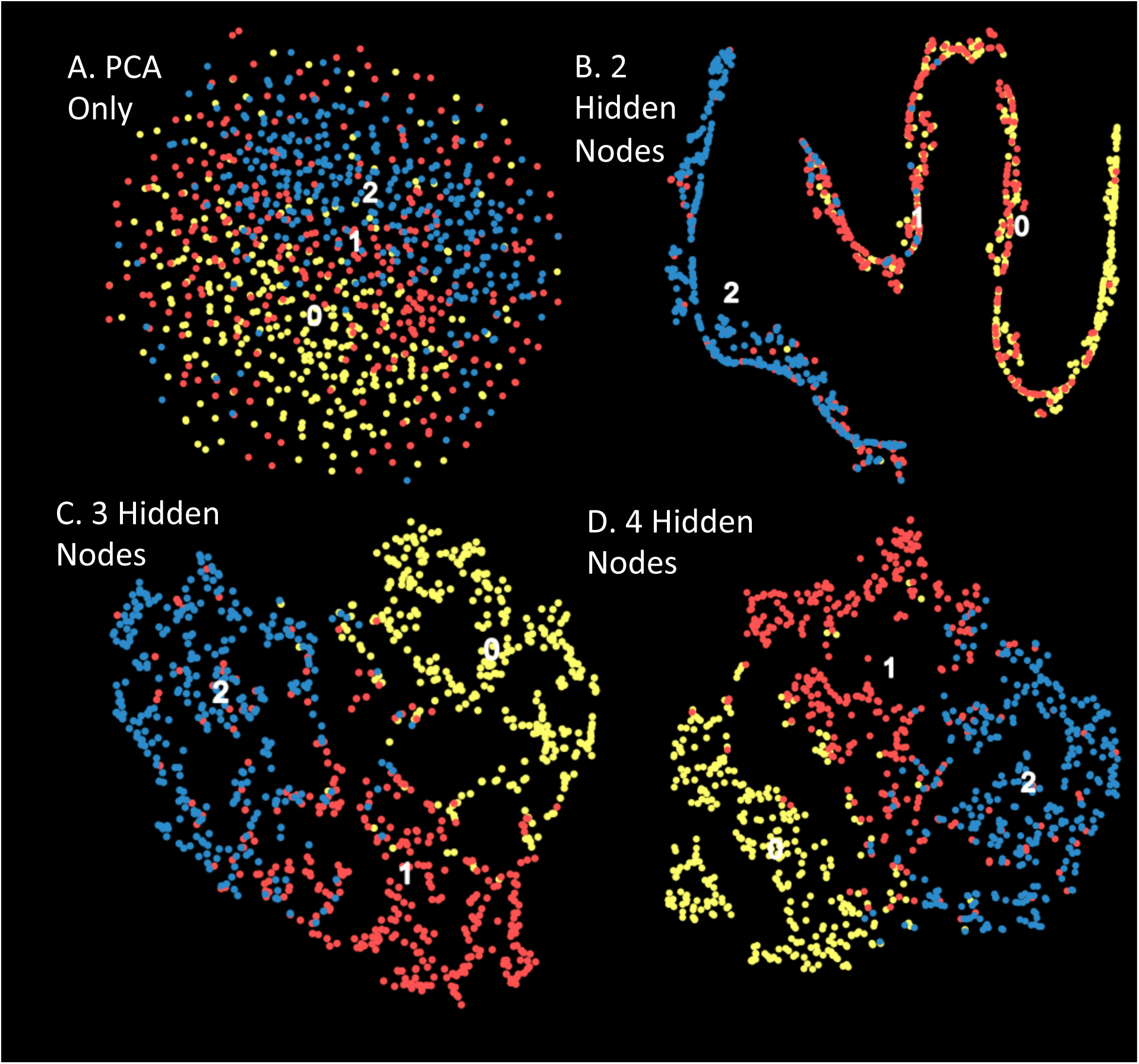
Case vs. Control clustering via principal components analysis and t-distributed stochastic neighbor embedding after training the DA for controls and cases generated from a combination of models 1 and 2. Controls are shown in yellow, subtype 1 (model 1) is shown in red, subtype 2 (model 2) is shown in blue. **A)**. Raw input. **B.)** 1,000 training epochs with 2 hidden nodes. **C.)** 1,000 training epochs with 3 hidden nodes. **D.)** 1,000 training epochs with 4 hidden nodes.

### ALS Survival Analysis

We evaluated the DAPS’ ability to predict ALS patient survival (Fig 6A) with ten-fold cross validation. Both models used a random forest regressor to predict survival to compare predictions between the raw imputed data and the hidden layer of a 250 node DA.

**Fig. 6.**
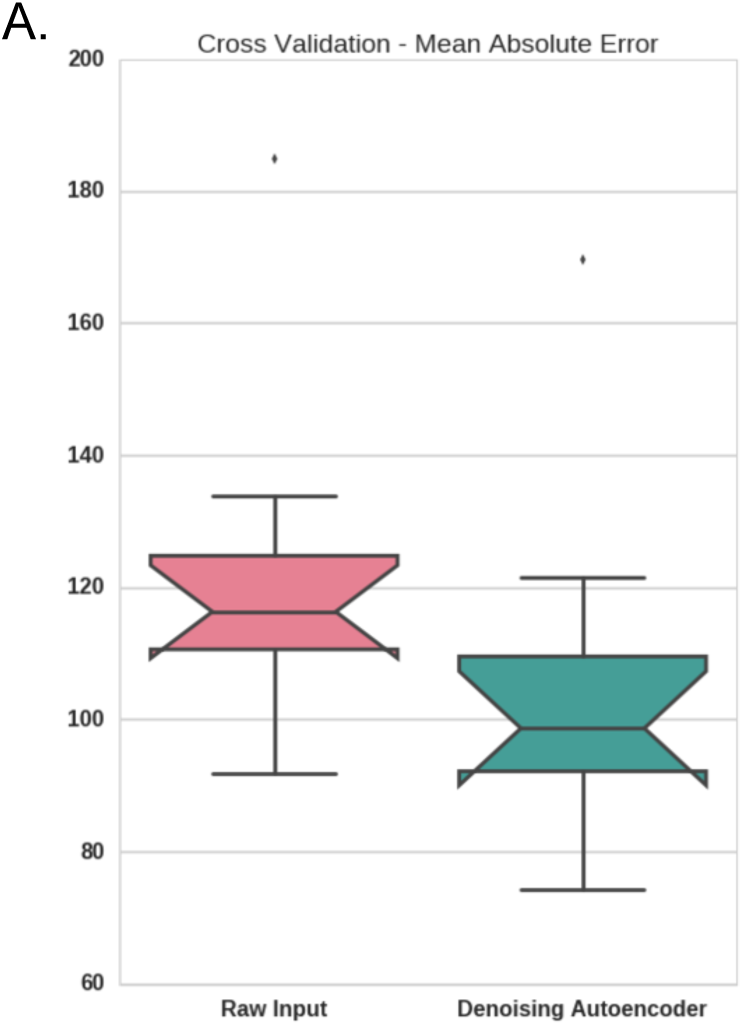
**A.)** Ten-fold cross validation survival prediction. Mean absolute error in days. Notch indicates 95% confidence interval for the median. Whiskers extend 1.5 times past the low and high quartiles. Points outside this range are denoted as dots and represent outliers.

Next we performed t-SNE clustering to compare PCA with the hidden layer of the DA (Fig 7 A-B, Supp. Fig. 7). The visualization constructed from the hidden layer of the DA shows several clear clusters with low patient survival as well as a more heterogeneous cluster with longer survival. PCA produced some patterns but did not produce any clear clustering.

**Fig.7.**
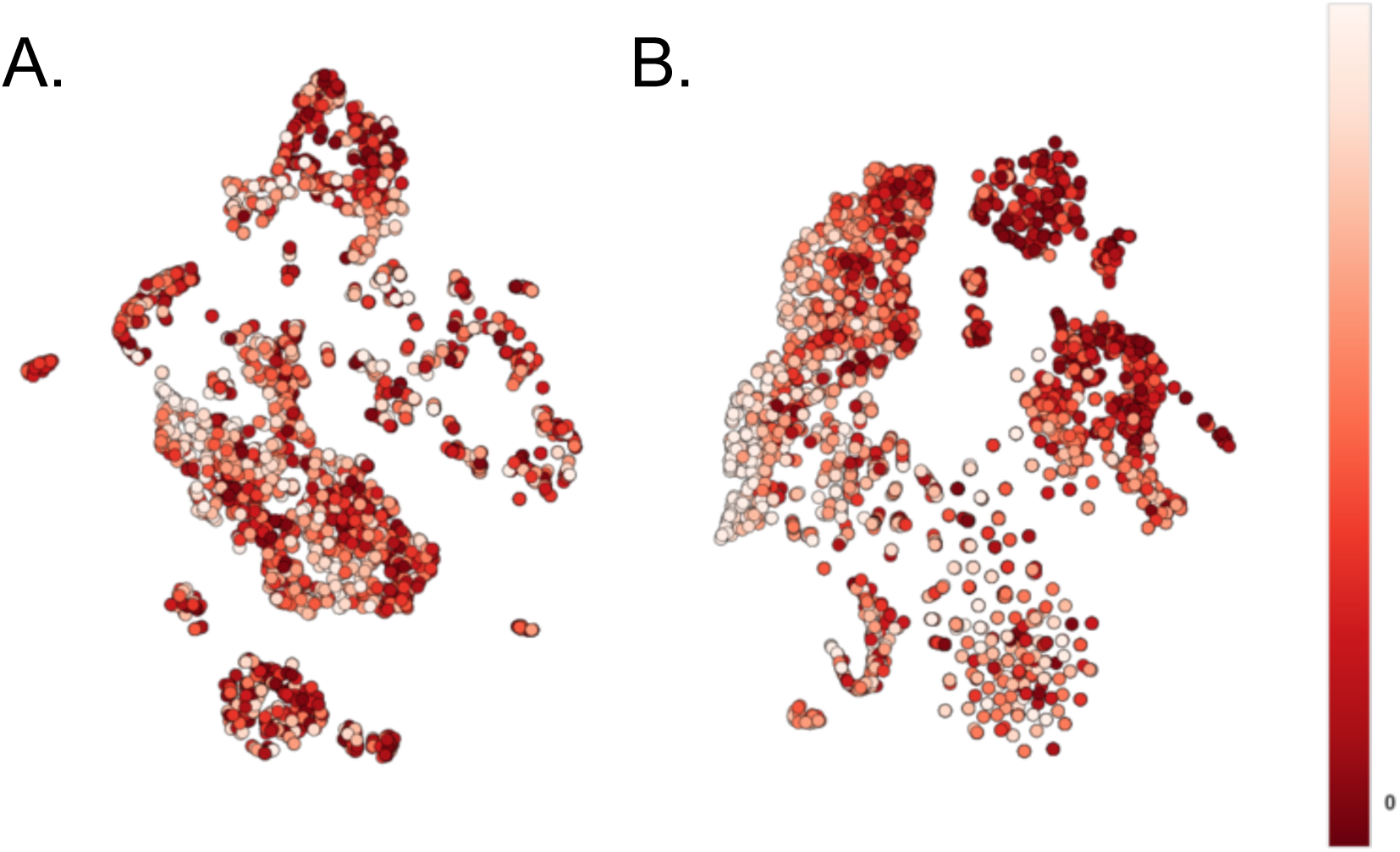
**A.)** PCA (2 components) followed by t-SNE. **B.)** t-SNE of the DA (250 nodes) hidden layer. All cluster coloring was determined by rank of days survived. Light colors indicate longer survival.

## DISCUSSION

In this study, we presented a semi-supervised learning approach using DAs to model patients in the EHR. The benefits of the method presented in this work are; 1) It is relatively inexpensive to perform analysis on large amounts of unlabeled EHR data; 2) It is expensive to have data labeled at a research quality by a clinician; 3) The DA+RF has strong performance when there are many unlabeled samples and few labeled samples.

Competitive supervised classification accuracy with a large degree of feature reduction indicates the DA successfully learned the structure of the high-dimensional EHR data. DAs are particularly well suited to the EHR because their unsupervised nature allows the formation of a semi-supervised classifier and the ability to utilize large un-annotated patient populations to improve classification accuracy. The dimensionality reduction of DAs allows clustering of the reduced feature set for the visualization and determination of subtypes. These clusters may reveal disease subtypes, fine-tuned targets for genotypephenotype association. The DA models are easily de-constructible because they use a simple model for the traditional classifier with transparent node compositions that can be traced back to inputs. In addition, our method proposes a straightforward modification to the DA to enable it to process missing data without imputation.

PheWASs are a powerful tool to leverage the vast clinical data contained in the electronic health record but currently suffer from the reliance on billing codes or manual clinician annotation. Denny et al. [1] call out the need for increased accuracy in phenotype definition in the original PheWAS publication, particularly for rare phenotypes or phenotypes that do not directly correspond with a billing code. In addition, several studies have found increased genetic linkage via subtyping [14–18,43]. Li et al. [19] presented a powerful example of EHR subtyping of patients with type 2 diabetes using a similar methodology, but they utilized Ayasdi, a commercial, closed source topology data analysis software tool. Our method is built on free, open source libraries that will continue to be improved and our software is accessible for the research community.

DA nodes and clusters of nodes provide composite variables that may better approximate and represent the condition of the subject. These additional phenotype targets may provide more homogeneous targets for genotype associations. Beyond genotype to phenotype association, these visualizations may also help clinicians to understand the level of heterogeneity for a specific disease and to make treatment associations among sub-clusters of patients. While further work is required to analyze the makeup and meaning of the ALS survival clusters recognized by DAPS, they suggest a helpful starting point for investigation.

Our work provides an important contribution but additional analysis and challenges remain. The transition from simulated data and relatively homogenous clinical trial data to diverse multi-disease real world clinical data will likely require additional steps. In addition, in our simulations we assume a preprocessing step has already been performed to handle the compound structure present in the EHR. This step is necessary to transform categorical, free text, images and temporal data to suitable input for the DA. The PROACT ALS clinical trial data does not currently include any free text or images.

Future work will focus on developing tools to examine and interpret constructed phenotypes (hidden nodes) and clusters. In addition, we will develop a framework for evaluating the significance of constructed clusters for genotype to phenotype association. We anticipate high weights indicate important contributors to node construction revealing relevant combinations of input features. Finally we will construct a scheme for determining optimal hyper parameter (i.e. hidden node count) selection.

## Acknowledgements

This work is supported by the Commonwealth Universal Research Enhancement (CURE) Program grant from the Pennsylvania Department of Health as well as the Gordon and Betty Moore Foundation's Data-Driven Discovery Initiative through Grant GBMF4552 to C.S.G. The authors would like to thank Dr. Jason H Moore for helpful discussions. The authors acknowledge the support of the NVIDIA Corporation for the donation of a TitanX GPU used for this research.

## SUPPLEMENTARY MATERIALS

### Parameter Sweep Specifications

**Supp. Table 1.** Simulation Model 2 Parameter Sweep Specifications.

| Parameter | Values |
| --- | --- |
| Observed Variables | 100 |
| Effect Magnitude (x Variance) | 5 |
| Hidden Input Effects | 1, 2, 4, 8 |
| Effectted Observed Variables per Hidden Input Effect | 5, 10 |
| Unlabeled Patients | 10,000 |
| Labeled Patients | 100, 200, 500, 1000, 2000 |
| Systematic Bias | 0.1 applied to 0.33 of patients |
| DA Hidden Nodes | 2, 4, 8 |

**Supp. Table 2.** Simulation Model 3 Parameter Sweep Specifications.

| Parameter | Values |
| --- | --- |
| Observed Variables | 100 |
| Effect Magnitude (x Variance) | 5 |
| Hidden Input Effects | 1, 2, 4, 8 |
| Effectted Observed Variables per Hidden Input Effect | 10 |
| Unlabeled Patients | 10,000 |
| Labeled Patients | 100, 200, 500, 1000, 2000 |
| Systematic Bias | 0.1 applied to 0.33 of patients |
| DA Hidden Nodes | 2, 4, 8 |

**Supp. Table 3.**
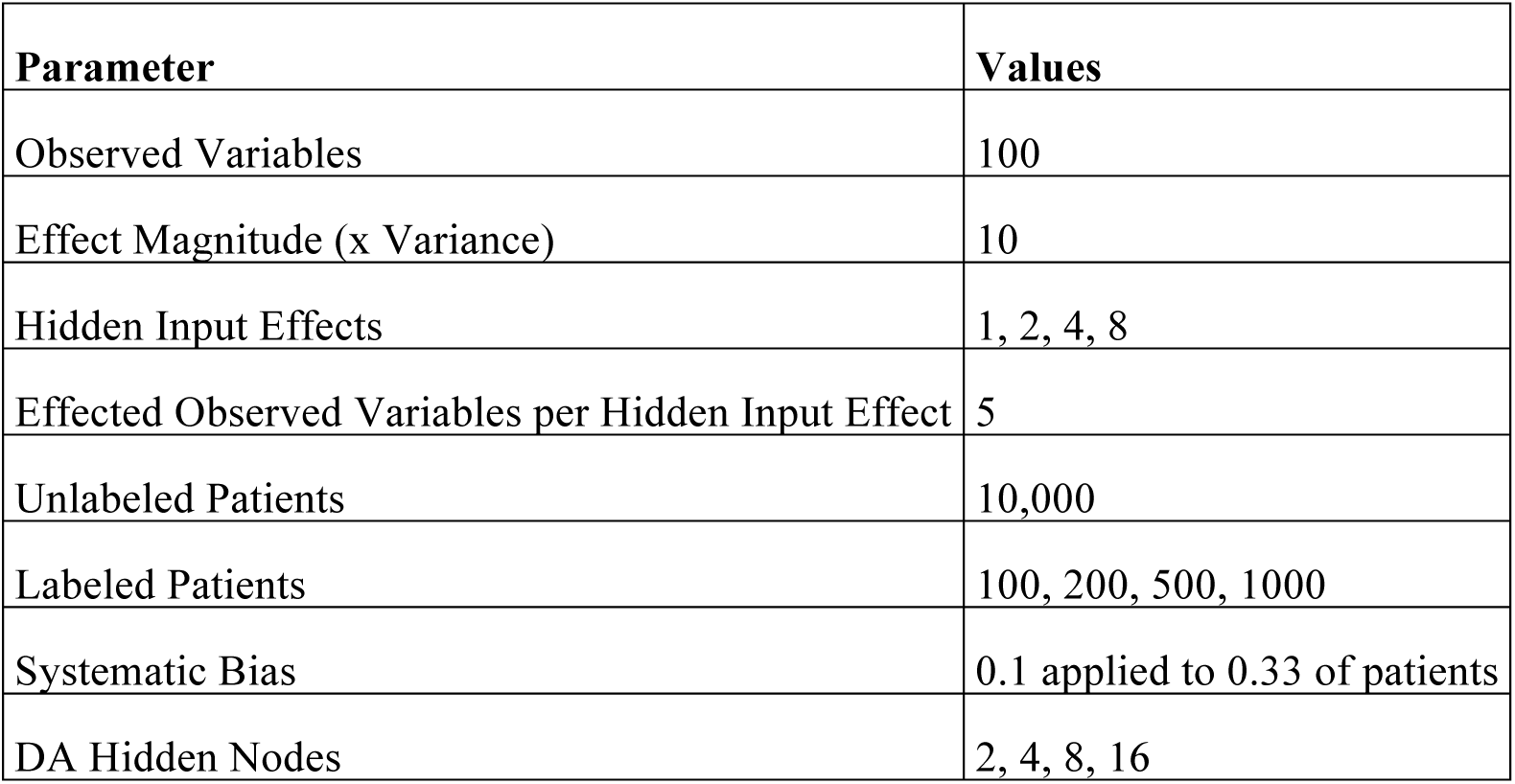
Simulation Model 4 Parameter Sweep Specifications.

**Supp. Table 4.** Missing Data, Simulation Model 1 Parameter Sweep Specifications.

| **Parameter** | **Values** |
| --- | --- |
| Observed Variables | 100 |
| Effect Magnitude (x Variance) | 2 |
| Hidden Input Effects | 4 |
| Effectuated Observed Variables per Hidden Input Effect | 10 |
| Unlabeled Patients | 10,000 |
| Labeled Patients | 200 |
| Systematic Bias | 0.1 applied to 0.33 of patients |
| DA Hidden Nodes | 2 |
| Missing Data | 0, 0.1, 0.2, 0.3, 0.4, 0.9 |

**Supp. Figure 1.**
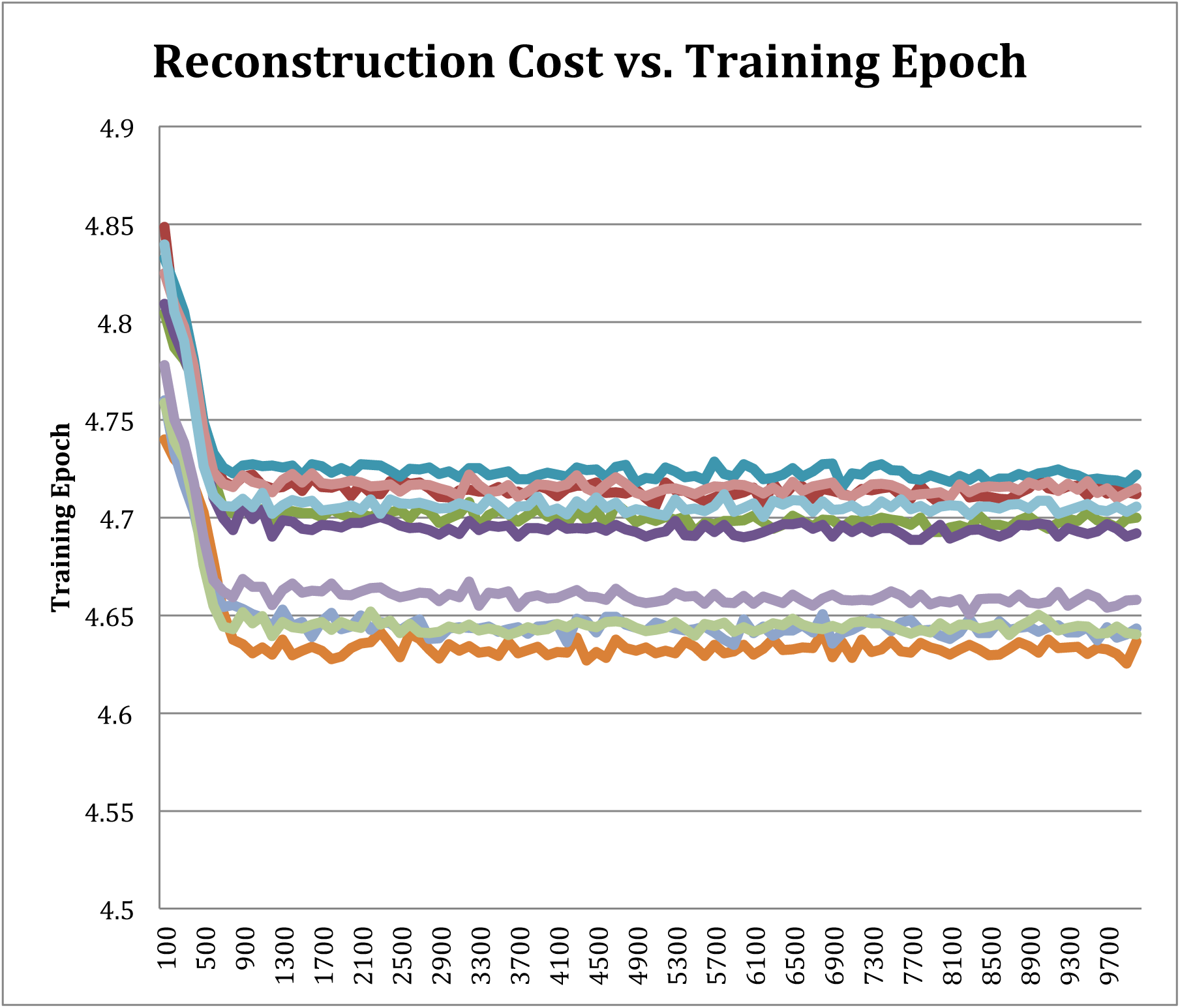
Denoising Autoencoder Reconstruction Cost vs. Training Epochs

**Supp. Figure 2.**
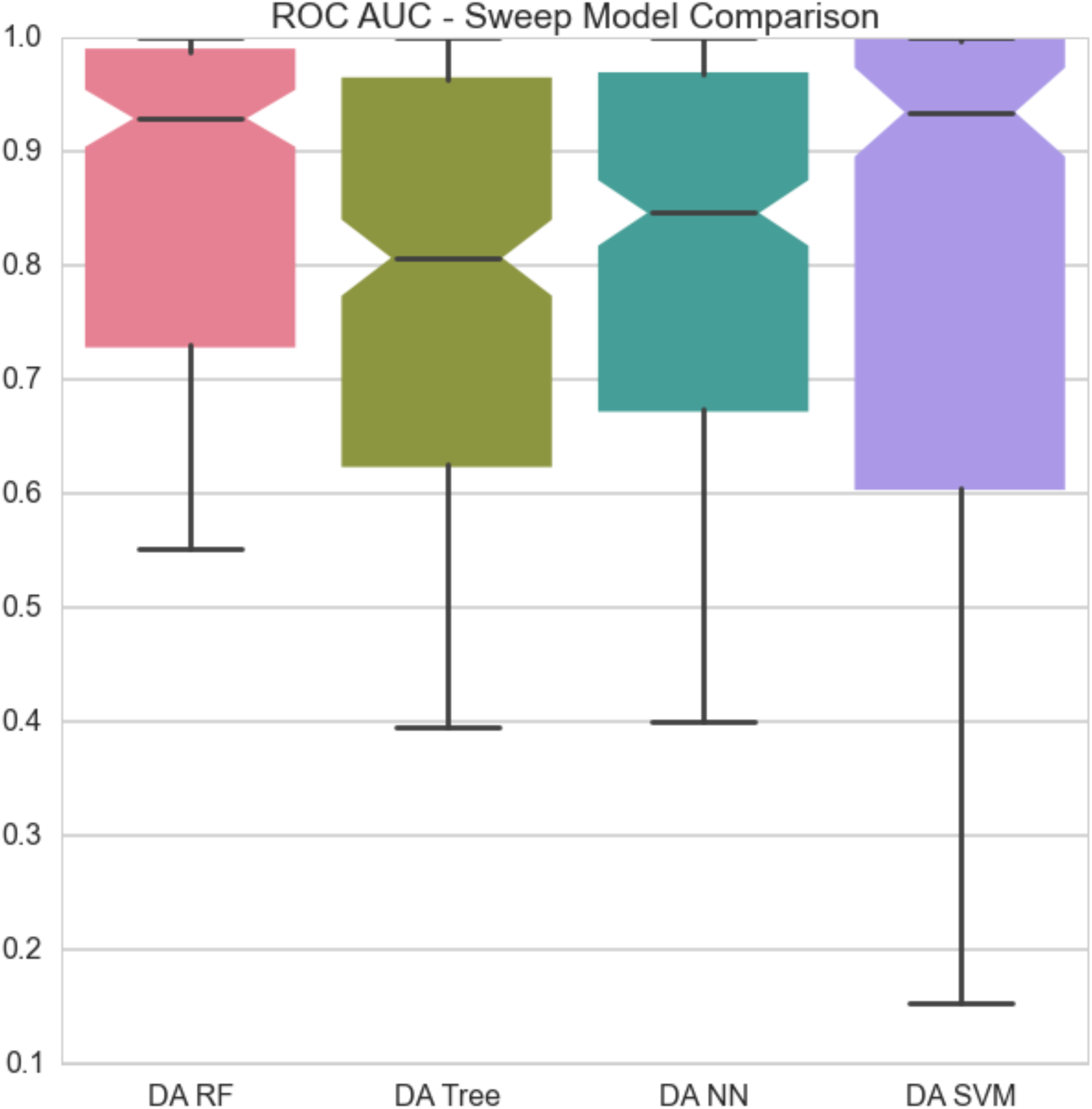
ROC AUC comparisons for traditional classifiers across model 1 with DA hidden nodes as inputs.

**Supp. Figure 3.**
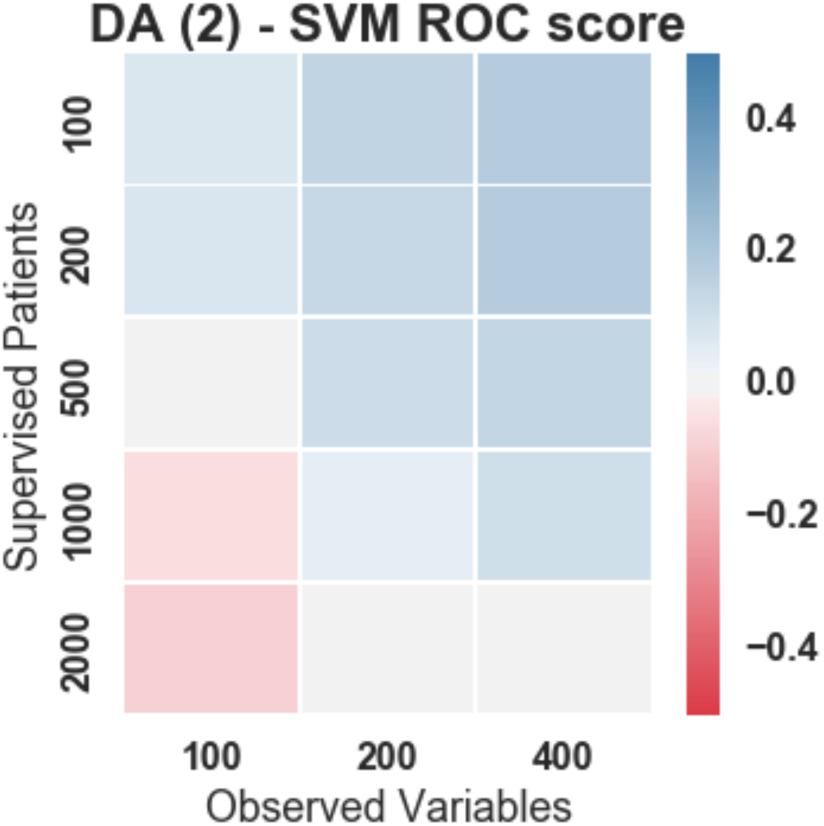
Heat Map showing the difference of DA+RFC and SVM methods in relation to the number of labeled patients and observed variables under simulation model 1.

**Supp. Figure 4.**
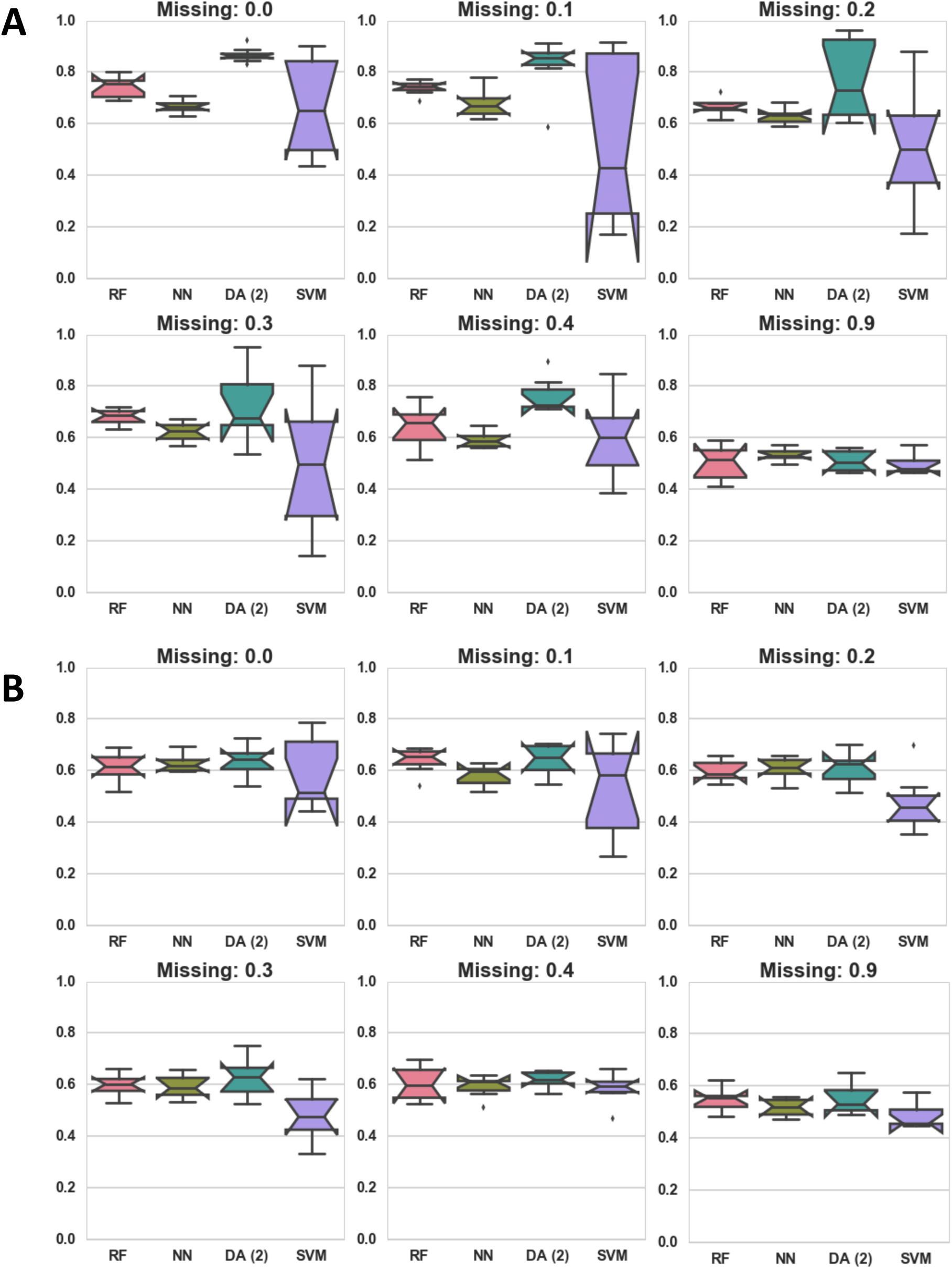
**A)** Classification AUC in relation to the amount of missing data under simulation model 1. **B)** Classification AUC in relation to the amount of missing data under simulation model 3.

**Supp. Figure 5.**
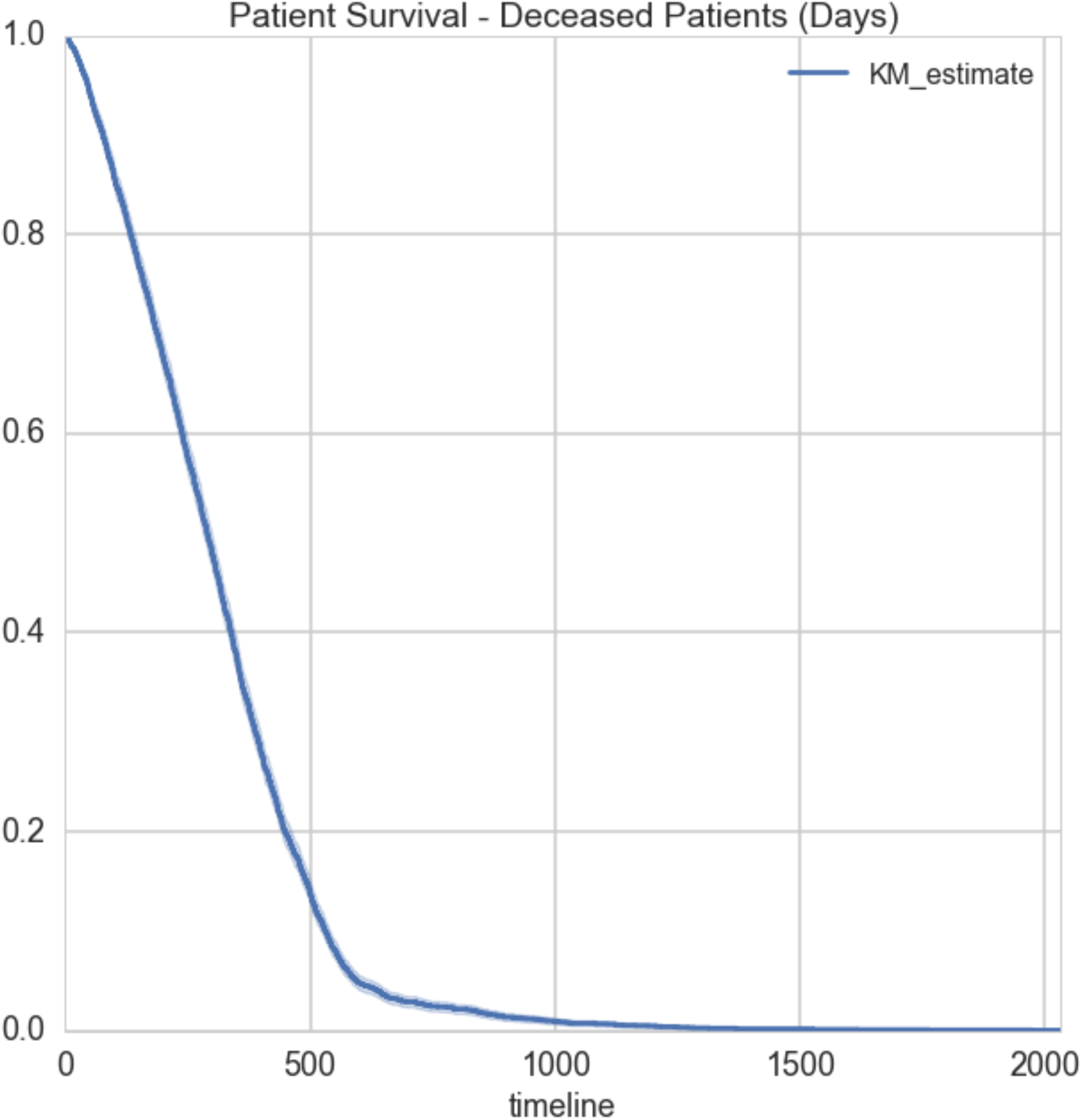
**A.)** Kaplan-meier curve showing patient survival in days. For this analysis only patients with known death dates were included.

**Supp. Figure 6.**
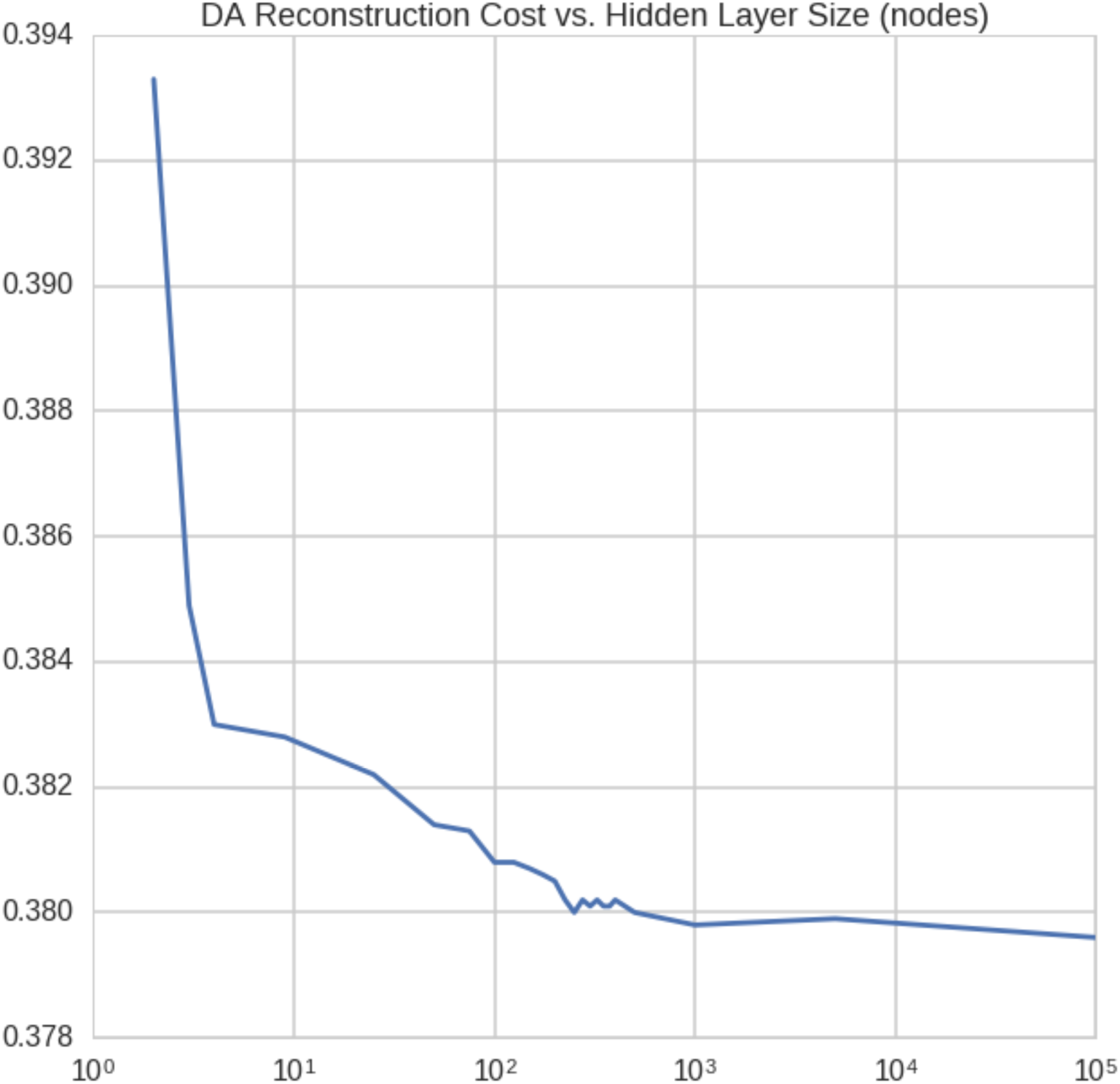
**A.)** Reconstruction cost in relation to the number of hidden nodes in the denoising autoencoder when trained on the PRO-ACT ALS data.

**Supp. Figure 7.**
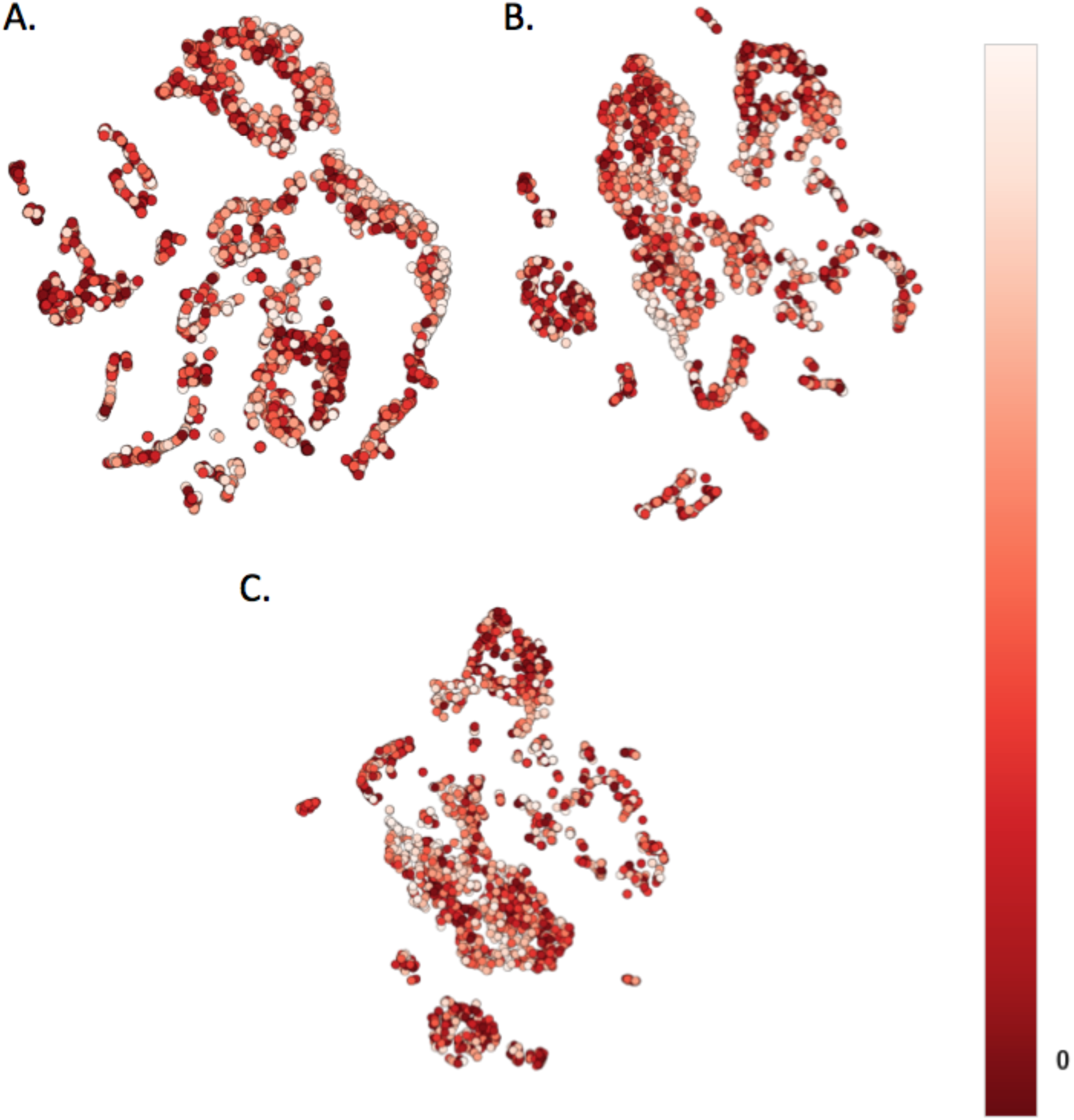
ALS patient clustering using Principal Components Analysis. Color indicates survival length, by ranking. Lighter colors survived longest. **A**) Four Principal Components. **B)** Eight Principal Components. **C)** Sixteen Principal Components.

